# Genotype grouping and bio-geographical analysis revealed the worldwide spatiotemporal spread of *Citrus yellow vein clearing virus*

**DOI:** 10.1101/2023.10.26.564128

**Authors:** Yongduo Sun, Raymond Yokomi

## Abstract

The *Citrus yellow vein clearing virus* (CYVCV) causes a viral disease that has been reported in specific citrus-growing regions in Euro-Asia including countries of Pakistan, India, Türkiye, Iran, China and south Korea. Recently, CYVCV was detected in a localized urban area in a town in heart of California’s citrus-growing region and marks the first occurrence of the virus in North America. CYVCV is spread by aphid and whitefly vectors and is graft and mechanically transmitted. Hence, it is an invasive disease that presents a significant threat to the California citrus industry, especially lemons which are highly susceptible to CYVCV. To elucidate the origin of the CYVCV California strain, we used long-read sequencing technology and obtained the complete genomes of three California CYVCV isolates, CA1, CA2, and CA3. The sequences of these isolates exhibited intergenomic similarities ranging from 95.4% to 97.4% to 54 publicly available CYVCV genome sequences which indicated a relatively low level of heterogeneity. However, CYVCV CA isolates formed a distinct clade from the other isolates when aligned against other CYVCV genomes and coat protein gene sequences. Based on a rooted Maximum Likelihood phylogenetic tree, CYVCV CA isolates shared the most recent common ancestor with isolates from India. Further examination of 79 coat protein gene sequences collected over a 31-year period that spanned regions from East and South Asia to the Middle East and California, Bayesian evolutionary inferences resulted in a spatiotemporal reconstruction that placed the origin of all CYVCV to the 1930s, with South Asia as the most plausible geographic source. This analysis also suggested that CYVCV CA isolates diverged from Indian lineages, possibly around the 2010s. Moreover, the spatiotemporal phylogenetic analysis indicated two additional virus diffusion pathways: one from South Asia to East Asia and another from South Asia to the Middle East. Collectively, our phylogenetic inferences offer insights into the probable dynamics of global CYVCV dissemination, emphasizing the need for citrus industries and regulatory agencies to closely monitor citrus commodities crossing state and international borders.

**Author Summary:** A localized outbreak of CYVCV was detected in a central California town, marking its first appearance in North America. The study sequenced the complete genomes of three CYVCV isolates from California and employed statistical algorithms to investigate the population dynamics and origin of CYVCV. Upon comparing coat protein gene sequences, the CYVCV isolates from California formed a distinct group separate from those found in other geological regions. The study’s spatiotemporal phylogenetic analysis highlighted that CYVCV likely originated in the 1930s, with South Asia as the most plausible source. Notably, the CYVCV isolates from California diverged from Indian lineages, possibly around the 2010s. This study contributes to a better understanding of CYVCV’s genetic and molecular diversity, shedding light on virus ecology, evolution, and biology.

## Introduction

The *Citrus yellow vein clearing virus* (CYVCV) presents a pressing quarantine concern regarding transport of citrus commodities and international trade. The manifestations of CYVCV disease exhibit significant variations contingent upon citrus varieties and prevailing environmental conditions (1). Lemon (*Citrus limon*) and sour orange (*C. aurantium*) trees are highly symptomatic, while a broad range of other citrus cultivars, though susceptible, remain asymptomatic (2, 3). CYVCV-symptomatic citrus trees display stunted growth, diminished citrus yields, yellow vein clearing, water-soaked appearance of veins on the adaxial side, leaf deformities, intermittent ringspots, and venial necrosis (1–3). No effective management strategies have been found to counteract the deleterious impact of CYVCV since the citrus host range is wide and the insect vectors are common in citrus orchards.

CYVCV is a member of the *Alphaflexiviridae* virus family, *Mandarivirus* genus, and constitutes a positive-sense flexuous RNA virus (4, 5). It is noteworthy that CYVCV shares a high genome similarity, of approximately 74% sequence identity with *Indian citrus ringspot virus,* another member of the *Mandarivirus* genus (6). To date, only a handful of CYVCV isolates have been subjected to complete sequencing (3, 7, 8). The viral genome encompasses approximately 7530 base pairs and encodes six predicted open reading frames (ORFs). ORF1 encodes a solitary polyprotein, with four constituent subunits, namely methyltransferase, oxygenase, RNA helicase, and RNA-dependent RNA polymerase. ORF2 to ORF6 encode triple gene block gene 1 (TGB1), TGB2, TGB3, coat protein (CP), and a nucleic acid-binding protein (1, 5). Our understanding of the functional properties of CYVCV-encoded proteins remains limited. CP has been identified as an RNA silencing suppressor (9) and has been linked to the severity of symptoms in citrus. Subsequent research has elucidated the CP’s interaction with the 40S ribosomal subunit protein S9-2, whose transient accumulation in the host impedes the CP’s silencing suppressor activity (10).

CYVCV can be transmitted through grafting and mechanical means and is naturally vectored by at least three aphids (*Aphis spiraecola, A. craccivora, and A. gossypii*) and the citrus whitefly, *Dialeurodes citri* (11, 12). The initial report of yellow vein clearing disease was in lemon and sour orange in Pakistan in 1988 (13). Since then, it has been reported in various locales, including Türkiye, India, Iran, China, and South Korea (2, 14–16). The rapid proliferation of CYVCV in China since 2009 has resulted in substantial losses in lemon production where the disease incidence with high (17). In 2022, during a routine multi-pest survey conducted by the California Department of Food and Agriculture (CDFA), CYVCV-infected citrus trees were identified in localized urban properties in the city of Tulare, California, United States of America (USA), although surveys of nearby citrus orchards indicate no spread yet to commercial citrus (7). The United States Department of Agriculture, Animal and Plant Health Inspection Service also tested samples from infected trees and verified the diagnosis of CYVCV. The CDFA currently designates CYVCV as a pest of high concern (Pest rating A).

The evolutionary dynamics of CYVCV remain enigmatic. This knowledge gap is, in part, attributable to the scarcity of CYVCV sequence data. Additionally, the virus’ evolutionary and ecological dynamics may have developed on different time scales, a phenomenon observed in other plant viruses (18). In a 2019 report, a phylogenetic tree constructed using the maximum likelihood method delineated CYVCV isolates into two clusters. CYVCV isolates from India, Türkiye, and Pakistan coalesced in one group, while those from China formed another (6). Recent developments, including CYVCV’s expansion into citrus-growing regions in California and South Korea, have drawn attention to the virus’s origin, global transmission, and spread (7). As more CYVCV genotypes are identified and CYVCV genotypes become better defined, opportunities for reanalyzing previously generated datasets to refine CYVCV genotype differentiation will emerge. Furthermore, new CYVCV genome sequences will be invaluable for comparative genomics, contributing to a deeper understanding of CYVCV’s etiology, relationships, and evolution. In this study, employing state-of-the-art long-read sequencing technology, we successfully obtained the complete genome sequences of three novel CYVCV California isolates. Through genotypic clustering and Bayesian evolutionary analysis, we elucidate the spatiotemporal and phylodynamics of CYVCV on a global scale.

## Results

### Field survey and CYVCV qPCR detection

A diverse collection of CYVCV was acquired from known infected trees (*C. limon, C. hystrix, C. reticulata, C. paradisi, Fortunella* sp.) from different properties in Tulare, California, USA and propagated in the greenhouse (Figure 1A). Graft propagations in sour orange (*C. aurantium* L.) and Eureka Lemon (*C. limon*) exhibited clear CYVCV symptoms of vein clearing, water soaking, and leaf distortion (Figure 1B); whereas virus propagations in mandarin (*C. reticulata*), *C. macrophylla*, Duncan grapefruit (*C. paradisi*), Madam Vinous sweet orange (*C. sinensis* (L.) Osbeck), and S1 citron (*C. medica* L.) remained asymptomatic (not shown). Systemic CYVCV infection was confirmed by RT-qPCR using United States Department of Agriculture-Animal and Plant Health Inspection Service-Plant Protection and Quarantine (USDA-APHIS-PPQ)-approved CYVCV primers. An intriguing observation emerged when analyzing Eureka lemon plants: virus titers were notably higher in flowers compared to stems and leaves, despite the absence of obvious growth defects or disease symptoms in flowers (Figure 1C).

**Fig 1.**
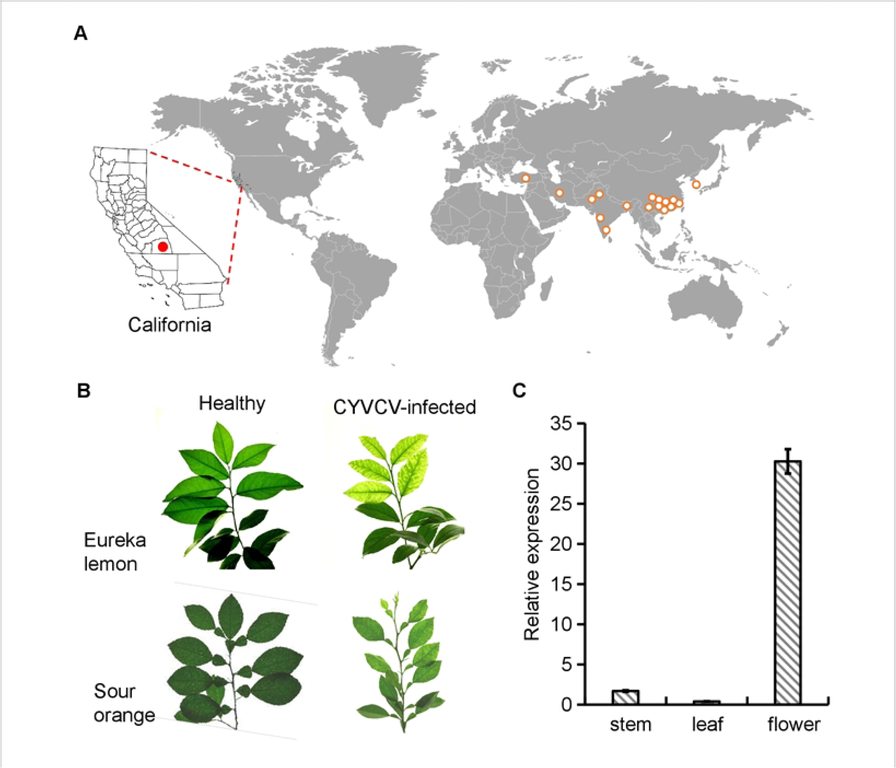
*Citrus yellow vein clearing virus* (CYVCV) outbreaks and traits. **A.** Global distribution of CYVCV. The global occurrence of CYVCV, except South Korea, was derived from data reported in the European and Mediterranean Plant Protection Organization Global Database, accessed in August 2023. The emerging of CYVCV in Tulare County, California, United States was marked with a red dot. All the others were marked with orange circles. **B.** A typical branch of Eureca lemon *(Citrus limon)* and Sour orange (*C. aurantium*) trees showing a typical CYVCV-induced yellow vein clearing phenotype in the greenhouse. C. RT-qPCR data shows the distribution patterns of CYVCV in lemon stem, leaf, and flower. The citrus *Nad5* gene was used as an internal control. Three biological test was repeated and a similar trend was obtained.

### The whole genome sequences of CYVCV CA isolates and nucleotide diversity analysis

Three CYVCV isolates were selected that represented different CYVCV sources propagated from six different infected field trees collected over a 5.2-sq km area of Tulare. The complete genomes of these three isolates were sequenced and designated as CYVCV CA1 (Accession number OR037276.1), CYVCV CA2 (Accession number OR670060), and CYVCV CA3 (Accession number OR670061). This addition of three new CYVCV California isolates brings the total number of reported CYVCV whole genome sequences to 57 (Table 1). These genomes of the three California isolates consisted of 7530 nucleotides (nt), excluding the 3’ poly A tail, and harbored six open reading frames (ORFs). It is noteworthy that the genome size of CYVCV isolates ranged from 7528 nt to 7531 nt due to insertion/deletion mutations at positions 20, 29, 30, and 6127 nt (Supporting Figure 1). Multiple sequence alignment displayed base-pair differences with variations, which indicated that CYVCV CA isolates exhibited a relative high divergence compared to the reference sequence CYVCV CQ isolate (Accession number NC_026592.1) isolate from China (Figure 2A). Sequence identity analysis unveiled that the reported global CYVCV isolates share a high sequence similarity, ranging from 95.1% to 100%, indicated a relatively low level of heterogeneity. (Figure 2B). Within this spectrum, the sequences of these CYVCV CA isolates exhibited intergenomic similarities ranging from 95.4% to 97.4% to 54 publicly available CYVCV genome sequences. In contrast, the CYVCV CA in-group isolates exhibited a relatively high sequence identity, reaching 99.6%. Notably, two pairs of CYVCV isolates share a hundred percent identity, as the CYVCV AY112 (Accession number MW429487.1) is identical to CYVCV AY204 (Accession number MG878869.1) and CYVCV CQ isolate NC_026592.1 is the same as another CYVCV CQ isolate KP313240.1.

**Fig 2.**
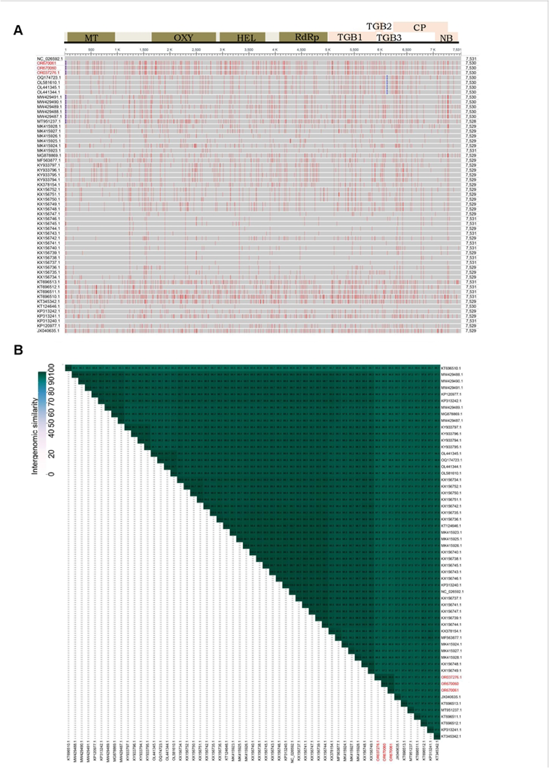
*Citrus yellow vein clearing virus* (CYVCV) nucleotide diversity. **A.** Multiple alignment reveals the base-pair with frequency-based difference, which was marked with red in the figure. **B.** VIRDIC generated heatmap incorporating intergenomic similarity values (right half) and alignment indicators (left half and top annotation).In the right half, the color-coding allows a rapid visualization of the intergenomic similarity of CYVCV genomes. The CYVCV CA isolates were marked with red.

### CYVCV genotype groups

Phylogenetic analysis using neighbor-net reconstruction of CYVCV complete genomes unveiled two major genotype groups (Figure 3A). All CYVCV isolates from China and South Korea formed a major group termed the "East Asia group," which further comprised eight subgroups, including South Korea subgroup (SK1) and China subgroup1-7 (C1-C7), named in chronological order of sample collection. Other CYVCV isolates were grouped into one major genotype, encompassing isolates from India, Pakistan, Türkiye, and California (USA). However, unlike the East Asia group, limited data availability hindered sub-clustering within this second major group. Nucleotide diversity analysis suggested closer genetic relationships between CYVCV isolates from India, Pakistan, Türkiye, and California.

**Fig 3.**
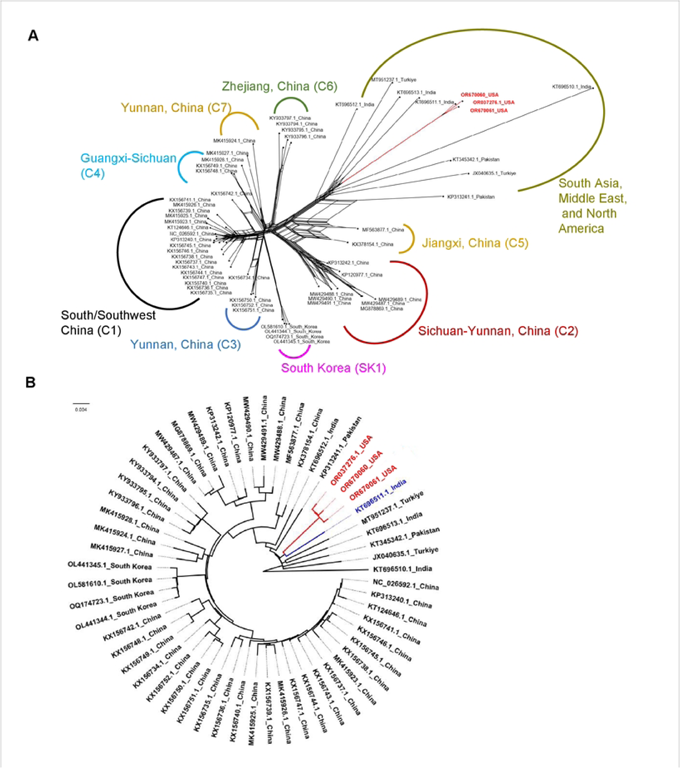
Construction of non-rooted and rooted phylogenetic tree upon *Citrus yellow vein clearing virus* (CYVCV). **A.** Neighbor network reconstruction of the complete genomes of 57 CYVCV isolates. The clade of CYVCV CA isolates are marked red. **B.** Maximum likelihood phylogenetic analysis of the sequences of 53 CYVCV isolates, excluding four isolates identified as recombinant. The clade of CYVCV CA isolates are marked with red. An India virus isolate in the same clade with CYVCV CA isolates is labeled with blue.

### Dissecting the origin of CYVCV CA isolates based on whole genome data

To ascertain the origin of CYVCV CA isolates, we initially assessed whether these isolates were the result of recombination. Among the 57 submitted CYVCV genome sequence data, four recombination events were detected. The CYVCV GX-STJ isolate (Accession number KX156742.1), CYVCV CQ-PO isolate (Accession number KX156735.1) CYVCV AY221 isolate (Accession number MW429491.1), and CYVCV PALI isolate (Accession number KT696512.1) were identified as recombinants. The analysis revealed no recombination events among CYVCV CA isolates (Supporting Table 1). Subsequently, a maximum likelihood phylogenetic analysis was performed using 53 CYVCV genome sequences, excluding the four identified as recombinants among the 57 submitted. This analysis also grouped CYVCV isolates into two major groups (Figure 3B). CYVCV isolates from China and South Korea clustered together, while isolates from other regions formed a separate group. Notably, CYVCV CA isolates shared the most recent common ancestor with an India CYVCV RMGI isolate (Accession number KT696511.1). This analysis suggested that CYVCV likely originated from India, with the India CYVCV ECAI isolate (Accession number KT696510.1) connecting directly to the root of the tree. Isolates from India, Pakistan, Türkiye, and the United States appeared to be more closely related to the ancestor of CYVCV compared to isolates from East Asia. It’s worth mentioning that no whole genome sequences were obtained from Iran, despite reports of CYVCV presence in 2007.

### Estimation of selection pressure of CYVCV-encoded ORF

To increase the dataset size, we evaluated the selection pressure of each CYVCV-encoded open reading frame (ORF) and sought a variable ORF for further analysis. The dN/dS ratio, indicating selection pressure, was calculated for all six coding regions. The results suggested that all six coding regions of CYVCV had evolved under varying degrees of purifying (negative) selection pressure (Supporting Table 2). The RdRP region exhibited the highest degree of purifying selection pressure (dN/dS=0.116), while the TGBs and CP regions displayed values ranging from 0.129 to 0.269. In contrast, the NB coding region exhibited high variability with a dN/dS ratio of 0.418, indicating the least purifying selection pressure. Given the negative selection pressure, all ORFs were presumed to be conserved. Considering the availability of more sequence data for the CP region compared to other ORFs and its relatively high purifying selection pressure, we selected the CP protein for further analysis.

### CYVCV grouping upon CP sequences

A total of 79 CP sequences were retrieved from GenBank, NCBI (Table 2). Neighbor-net analysis, based on CP sequences, classified CYVCV isolates into four major groups: East Asia (China and South Korea), Middle East (Iran and Türkiye), South Asia (India and Pakistan), and North America (United States). These groups were named according to their geographical distribution (Figure 4A). Notably, CYVCV CA isolates occupied a distinct clade separate from all other known isolates. This observation was further supported by a Direct Tissue Blot Immunoassay using a CP protein monoclonal antibody from CYVCV CQ isolate in China, which failed to detect the CP of CYVCV CA isolates (Supporting Figure 2). A maximum likelihood rooted tree revealed that CYVCV CA isolates/North America groups shared the most recent common ancestor with several India isolates (Accession numbers: KT696516.1, KT696518.1, KT696520.1, AOO32386.1) (Figure 4B). Thus, consistent with results obtained from whole genome sequences, phylogenetic analysis based on CP sequences suggested that CYVCV CA isolates may have originated from India.

**Fig 4.**
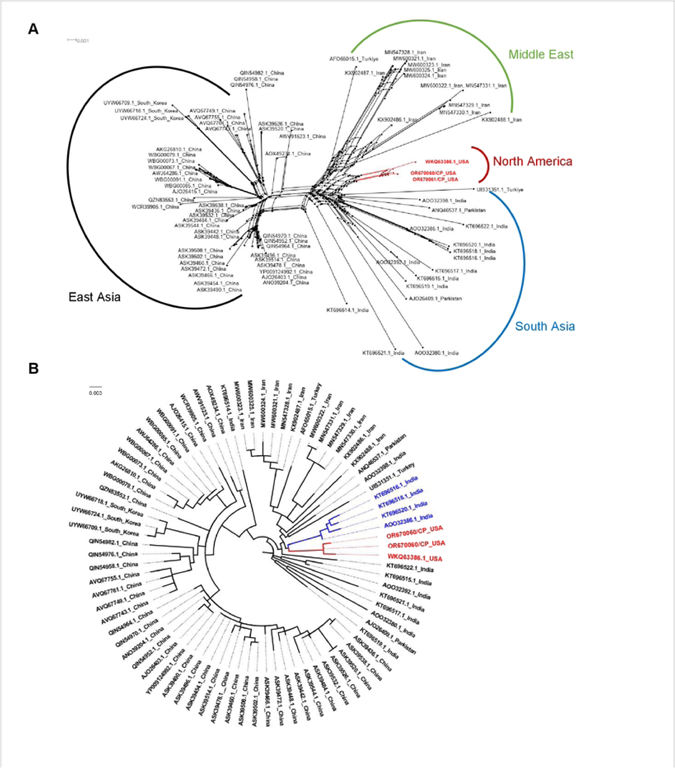
Construction of non-rooted phylogenetic tree upon *Citrus yellow vein clearing virus* (CYVCV) coat protein (CP) sequence. **A.** Neighbor network reconstruction of the CP sequence of 79 CYVCV isolates. The clade of CYVCV CA isolates are marked red. **B.** Construction of a Maximam likelihood tree to trace the most recent ancestor of CYVCV CA isolates upon the 79 CP sequence examinated in this study.The clade of CYVCV CA isolates are marked red. Four India virus isolates in the same clade with CYVCV CA isolates are labeled with blue.

### Reconstructing the global spatiotemporal transmission of CYVCV

To investigate the biogeographical diffusion patterns of CYVCV, we employed a Bayesian phylodynamic framework using the coat protein gene sequences collected at various time points. Root-to-tip analysis indicated weak temporal structure in the dataset, with the correlation coefficient equals to 0.2457 (Supporting Figure 3). Maximum clade credibility (BBM) tree from Bayesian molecular clock analysis estimated the origin of CYVCV from its progenitor to be over 93 years ago, although the first report of CYVCV was in 1988 (Figure 5). This finding implies that the CYVCV population may circulate for many years in a hypothetically restricted region before the global transmission. The most recent common ancestor of the CYVCV CA lineage appeared around 2010, 12 years prior to its documented discovery in California. Additionally, the CYVCV CA lineage shared a common ancestor with an India lineage, suggesting an origin around the 1950s (Figure 5). Thus, in line with geographical distribution and historical records, it is plausible that CYVCV CA isolates originated in South Asia, potentially India.

**Fig 5.**
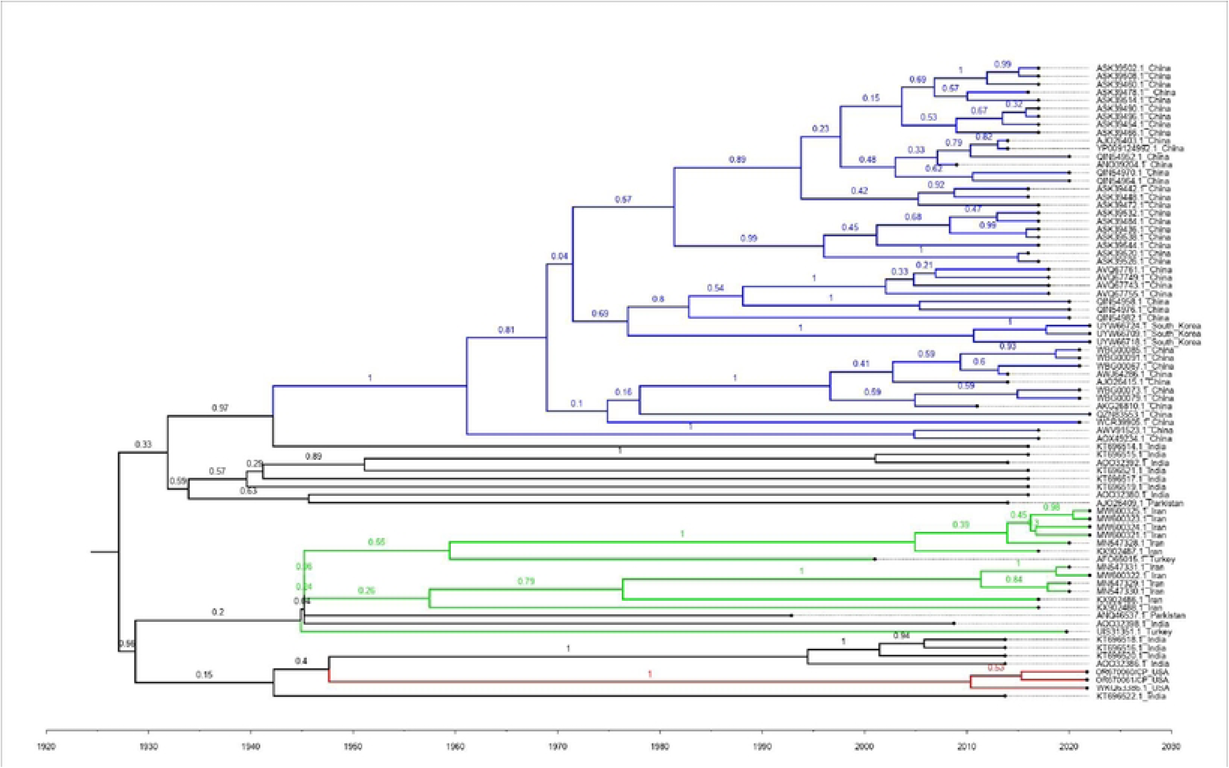
Maximum clade credibility BEAST tree reconstructed upon the 79 *Citrus yellow vein clearing virus* coat protein sequences The different color of the clade indicates different geological distribution: Blue: East Asia (China and South Korea); Green: Middle East (Iran and Turkiye); Red: North America (United States); Black: South Asia (India and Parkistan). Branch labels display posterior probabilites. The tree was visualized and modified with FigTree v.1.4.3.

Next, a phylogenetic tree showing the ancestral distribution ranges based on the BBM model was reconstructed upon program Reconstruct Ancestral States in Phylogenies (RASP) (Figure 6A) and, subsequently, a virus global diffusion map was suggested (Figure 6B). In line with the previous suggestion, the most likely distribution region of the most recent common ancestor of CYVCV CA isolates locates in South Asia. Moreover, the spatiotemporal phylogenetic analysis unveiled two additional virus diffusion pathways: one from South Asia to East Asia and another from South Asia to the Middle East. The common ancestor of Middle East isolates was inferred to have emerged around the 1950s, while the common ancestor of East Asia isolates arose after the 1960s. CYVCV GJ isolates from South Korea were suggested to have originated from CYVCV isolates in Yunnan, China (Supporting Figure 4).

**Fig 6.**
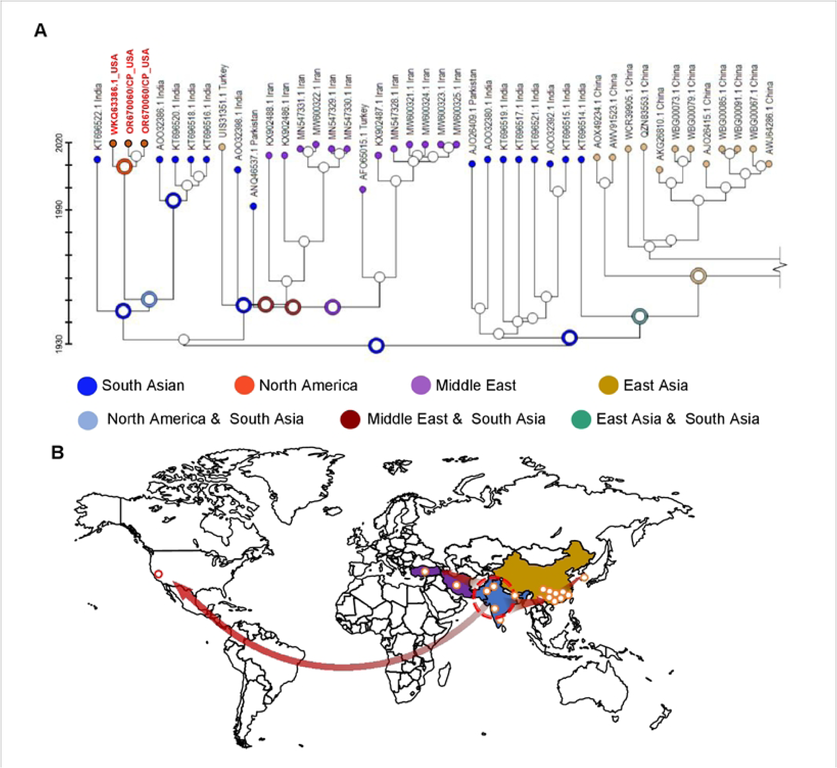
Time-calibrated biogeographic phylogenetic tree of *Citrus yellow vein clearing virus* (CYVCV). **A.** Cropped recostruction of CYVCV evolutionary history and dynamics of upon Bayesian evolutionary analysis. Corlored dots indicates the biogeographic distribution of CYVCV isolates. The circles illustrates the mostly likely distribution status of the most recent common ancestor in key nodes only. **B.** A schematic map exhibits the CYVCV outbreaks and hypothesized transmission routes. The sampling localities (colored spots) were catagoried into four regions of major clades, as North America shaded in orange, South Asia shaded in blue, Middle East shaded in purple, and East Asia shaded in sand, corresponding to those in the coat protein sequence neighbor network. The red arrows points to the referred transmission routes.

## Discussion

CYVCV, as an emerging viral disease in citrus, has become a major concern for some citrus-producing regions worldwide. It’s devastating effects, with yield losses of up to 50-80%, underscore the economic impact it carries (19). Over the past few decades, CYVCV isolates have been reported in seven countries: India, Pakistan, China, South Korea, Iran, Türkiye, and most recently, in California, USA (2, 14–16). Although the first observations of CYVCV-infected trees date back to 1988 in Pakistan, the first complete genome sequencing of CYVCV was not reported until four years later (13). In the case of Iran, CYVCV was reported in 2017, but to date, no whole genome sequences of CYVCV isolates from Iran have been made publicly available (16). The incursion of CYVCV into California, USA was documented in 2022; however, the genome sequence and molecular characteristics of CYVCV CA isolates have not been documented until this report. Our study leveraged cutting-edge long-read sequencing technology to obtain and annotate three CYVCV CA isolates from Tulare County, CA, to expand the CYVCV dataset and deepening our understanding of virus divergence which may become useful for disease management purposes.

The complete genome sequences of CYVCV CA isolates exhibit the typical genome organization of the *Mandarivirus* genus, ranging from 7,529 to 7,531 nt, excluding the 3′-poly (A) tail (1). The three CYVCV isolates obtained in this study encompassed 7,530 nt each. Global CYVCV isolates, when subjected to genome-wide comparison, displayed a high sequence similarity, ranging from 95.1% to 100%, indicating a limited degree of heterogeneity. Typically, the RdRP region of RNA viruses harbors functional domains of replicase proteins and is prone to nucleotide variability. The absence of proofreading activity in RNA polymerases of RNA viruses presents the potential for rapid evolution, genetic variability, and adaptation to new environmental conditions due to high mutation rates, resulting in the generation of variable populations (6, 20). In contrast, all six ORFs encoded by CYVCV did not exhibit significant variability, with an estimated selection pressure (dN/dS) below 1. The mechanism behind this observation requires further exploration, as it may shed light on the replication mechanism of CYVCV.

Studying the genetic and molecular diversity of viral pathogens contributes to a deeper understanding of virus ecology, evolution, and biology. In this context, our study aimed to investigate recombination and population dynamics using various statistical algorithms. The data obtained in this study revealed low levels of genetic diversity yet supporting the evolutionary relationships within the virus population. Neighbor network analysis, considering full genome sequences of CYVCV isolates from different countries, indicated that CYVCV isolates from India, Pakistan, and Türkiye were more closely related than those from China and South Korea. However, due to minimal nucleotide and amino acid differences, these isolates failed to form distinct clusters. The CP region of RNA viruses is vital for species demarcation, assessing genetic diversity, and developing immunodiagnostics (21, 22). Furthermore, only partial CP sequences are available for CYVCV isolates from Iran. Notably, a similar outcome was obtained. Analysis of CP sequences revealed that CYVCV isolates could be categorized into groups based on their geographic distribution, including South Asia, East Asia, Middle East, and North America. Notably, North America, at present, only encompasses the three isolates identified in this study. As the CYVCV genome pool expands, the boundaries of CYVCV genotypes will likely become more defined.

Maximum likelihood analysis of CP sequences indicated that CYVCV CA isolates diverged from isolates in South Asia, a region that has reported a substantial number of different CYVCV isolate sequences. In alignment with this observation, although CP gene sequences are generally conserved, the CP monoclonal antibody 18H5 of CYVCV CQ isolate from China, East Asia, did not effectively interact with CYVCV CA isolates. A previous CP analysis revealed that viruses in the *Mandarivirus* genus shared a common structural core and evolutionary origin (23). It is possible that divergent amino acid sequences in CYVCV CA isolates play a crucial role in the binding site with the 18H5 antibody. The minimal epitopes that could be recognized by the monoclonal antibody have not been identified. Further investigation is warranted.

In addition to elucidating the spatiotemporal scale of plant virus evolution, molecular sequence analyses can explore spatial population structure and provide insights into the transmission dynamics responsible for the current geographic distribution of plant viral lineages. Therefore, it is not surprising that plant virus epidemiology has started to incorporate statistical inference methods that combine temporal and spatial dynamics in a phylogenetic context (22, 24). For example, Bayesian phylogeographic methods have been applied to reconstruct the spatiotemporal history of *Tomato yellow leaf curl virus* spread and diversification. This analysis suggested that while the virus likely originated in the Middle East during the first half of the 20th century (25). Another example is *Cassava mosaic-like virus*, responsible for severe crop losses in sub-Saharan Africa, was estimated to have originated in mainland Africa in the late 1930s, with subsequent introductions to the southwest Indian ocean islands between 1988 and 2009 (26). Similarly, *Maize streak virus* (MSV), which has caused severe epidemics in maize-growing regions of Africa. Bayesian spatiotemporal reconstructions indicated southern Africa as the most probable origin of MSV at the beginning of the 20th century (27). In this study, Bayesian evolutionary analysis based on CYVCV CP sequences suggested that CYVCV CA isolates may have originated from South Asia, specifically India, around 2010. However, it is essential to recognize that spatiotemporal phylogenetic analysis has its limitations. First, the CYVCV isolate dataset remains relatively shallow, particularly in regions such as South India, Middle East, and North America. Additionally, the detection of plant viruses in perennial hosts is often delayed, as it takes time for symptoms to manifest. Different symptoms in different citrus varieties can further complicate disease diagnosis for citrus growers. For instance, while CYVCV is generally associated with vein clearing symptoms in sensitive cultivars, it can also produce ringspot symptoms on Kinnow mandarin and sweet orange, similar to those caused by ICRSV. Delayed detection in some commercial citrus varieties in India with ringspot symptoms was due to a lack of information about CYVCV’s ability to cause such symptoms (2, 15, 17). Therefore, caution must be exercised when interpreting phylogenetic relationships during a plant virus outbreak. Incorporating additional characteristics to support phylogenetic interpretation will likely yield more reliable inferences (28). For instance, it has been suggested that the host ecology determines the dispersal patterns of rice yellow mottle virus (18).

## Materials and Methods

### Sample collection, RNA extraction and real time-quantitative Polymerase Chain Reaction (RT-qPCR)

Citrus budstick samples from known CYVCV-positive citrus trees from the city of Tulare, California, were collected and propagated in a containment greenhouse in a variety of citrus cultivars. CYVCV source cultivars included Eureka lemon, mandarin, red grapefruit, kumquat, and makrut lime (*Citrus hystrix*). The grafted plants were maintained in an air-conditioned greenhouse at the San Joaquin Valley Agricultural Sciences Center in Parlier, California. The original host of CYVCV CA1 and CA2 was Eureka lemon while CA3 was from makrut lime. Total RNA from CYVCV the isolates were as extracted by Trizol Reagent (ThermoFisher Scientific, MA, USA) from leaves exhibiting symptoms of CYVCV.

A duplex RT-qPCR for simultaneous detection of CYVCV and the citrus *Nad5* gene as an internal quality control for nucleic acid extraction was employed. Specifically, RT-qPCR reaction was in a 10 µl reaction volume composed of 2 µl of RNA template, 5 µl of 2X reaction buffer, 0.4 µl each of CYVCV forward primer (5’-AAA TCC ATT AAC ACA GTG ACC TTC C-3’) and reverse primer (5’-AAC TCC TGA CAG TGC TCC AA-3’), 0.1µM of a CYVCV-specific 6-FAM/BHQ-1 labeled TaqMan probe (5’d FAM-CGTCGTTGCCAAGACACGCCA-BHQ-1), 0.4 µl each of Nad5 forward primer (5’-GATGCTTCTTGGGGCTTCTTKTT-3’) and reverse primer (5’-ACATAAATCGAGGGCTATGCGGATC-3’), and 0.1µM of a Nad5-specific VIC/QSY labeled TaqMan probe (5’d VIC-CAT AAG TAG CTT GGT CCA TCT TTA TTCCAT-QSY), along with 0.2 µl of iScript advanced reverse transcriptase and 0.9 µl of double-distilled water. This mixture was placed into a PCR plate, with cycling conditions encompassing reverse transcription at 50°C for 5 minutes, initial denaturation at 94°C for 2 minutes, followed by 40 cycles of denaturation at 94°C for 10 seconds and annealing/extension at 60°C for 40 seconds. RNA samples at a concentration of 10 ng/µl were tested in triplicate.

### CYVCV genome sequencing

A conserved region at the 5′ end, identified through alignment with other reported CYVCV isolates, served as the basis for designing a virus-specific 5’ race primer (5’-GGTTAGTGGTATTGCCCTGTT-3’). As for the 3’ race-specific primer, an oligo(dT) primer was employed. The amplicons generated from the 5’ and 3’ race PCRs were subjected to purification and subsequent cloning into the pGEM-T easy vector (Promega Corp., WI, USA). The constructed vectors were sequenced (Plasmidsaurus, OR, USA) to obtain the sequences of the CYVCV 5′ and 3′ termini.

Using these 5′ and 3′ termini sequences, the complete genome sequences were amplified for CYVCV CA isolates using the Q5 high-fidelity enzyme (New England Biolabs Inc., MA, USA) and virus-specific PCR primers (5′ primer-GAAAAGCAAACATAACCAACACACACCC; 3′ primer-CAGAAAATGGAAACTGAAAGCCTGAATATTT). This yielded a 7.5 Kb PCR amplicon which was sequenced with the latest long-read sequencing technology from Oxford Nanopore Technologies (ONT, Plasmidsaurus, OR, USA). The fully assembled genome sequences were annotated and deposited in GenBank under the accession numbers OR037276.1 (CYVCV CA1), OR670060 (CYVCV CA2), and OR6700601 (CYVCV CA3).

### Direct Tissue Blot Immuoassay (DTBIA)

The DTBIA was conducted in accordance with the procedure outlined in a report by (11). Briefly, young stems from CYVCV-infected lemon trees and healthy control samples were imprinted on nitrocellulose membranes. These membranes were then treated in 0.01 M PBS and subjected to blocking with 5% non-fat milk. The CYVCV monoclonal antibody 18H5 served as the primary antibody, while a secondary antibody comprising goat anti-mouse IgG (H&L) conjugated with alkaline phosphatase (ThermoFisher Scientific, MA, USA) was employed. The colorimetric signal was subsequently detected using nitro blue tetrazolium/5-bromo-4-chloro-3-indolyl-phosphate (NBT/BCIP, Sigma-Aldrich, MO, USA) at 37 °C for a duration of 10 minutes.

### Nucleotide diversity analysis

To analyze nucleotide diversity, the complete genomes of 57 CYVCV isolates were downloaded from the NCBI Virus Database and aligned using the NCBI Multiple Sequence Alignment Viewer. The Virus Intergenomic Distance Calculator (VIRIDIC) was employed to generate a heatmap utilizing default settings that incorporated intergenomic similarity values and alignment indicators (29).

### Construction of non-rooted phylogenetic neighbor network tree and rooted maximum likelihood tree

The complete genomes of 57 CYVCV isolates and 79 coat protein gene sequences (either full or partial gene sequences) were acquired from GenBank, NCBI, and aligned using MEGA 11 with the MUSCLE algorithm. The construction of a neighbor network was executed and subsequently modified utilizing SplitsTree 4, with 1000 bootstrap replicates (30). A maximum likelihood tree was generated using IQ-Tree (31). The maximum likelihood midpoint root tree was visualized and adjusted using FigTree v1.4.4.

### Recombination and estimation of selection pressure

An analysis of recombination between different CYVCV isolates was conducted using the Recombination Detection Program v4.56 (RDP4) software (32). The software utilized various algorithms, including RDP, GENECONV, CHIMAERA, MAXCHI, BOOTSCAN, SISCAN, and 3SEQ, each of which identified putative recombination events, major and minor parents, and breakpoints. Recombination events detected by at least four different methods were considered.

For estimation of selection pressure at the molecular level, the sequences of coding regions were aligned and analyzed according to Datamonkey (33). The rate of non-synonymous substitutions per nonsynonymous site (dN) to synonymous substitutions per synonymous site (dS) was analyzed separately for positive, negative, and neutral selection. Selection pressure was categorized as negative or purifying (dN/dS < 1), neutral (dN/dS = 1), and positive or adaptive selection (dN/dS > 1) for each dataset. To determine site-specific selection pressures acting on different genes, the codon-based likelihood algorithms Single-Likelihood Ancestor Counting (SLAC) were employed.

### Bayesian evolutionary inference

To assess the degree of divergence signal accumulated over the sampling time interval, the CYVCV CP sequence data was used, and an exploratory linear regression approach was conducted. Initially, a maximum likelihood (ML) tree was estimated under a non-clock (unconstrained) generalized time-reversible (GTR) + gamma substitution model using IQ-Tree (34). Root-to-tip divergences were plotted as a function of sampling time, employing a root maximized to yield the Pearson product moment correlation coefficient through TempEst (formerly known as Path-O-Gen) (35).

Subsequently, a time-calibrated phylogenetic tree was reconstructed using a Bayesian statistical framework from the software package BEAST v1.10.4 (24).The substitution model employed was HKY+ Gamma, and the clock type was set as strict clock. The length of Markov chain Monte Carlo (MCMC) chain was set as 20,000,000. Different tree priors, including Expansion growth, Exponential growth, Constant size, and Logistic growth, were tested and a Logistic growth tree providing the best fit was developed. BEAST employed MCMC integration to average over tree space, weighting each tree proportionally to its posterior probability. MCMC chains were visually checked by Tracer v1.6 and posterior parameters from tree samples were summarized via Treeannotator. A consensus tree was visualized and modified with FigTree v.1.4.3.

### Biogeographic analyses

Ancestral geographic ranges at each node were reconstructed using Statistical-Dispersal Vicariance Analysis (S-DIVA) and Bayesian Binary MCMC (BBM) analysis through the program Reconstruct Ancestral States in Phylogenies (RASP) (36). Four distribution ranges were defined based on geographic proximity as South Asia (India and Pakistan), East Asia (China and South Korea), Middle East (Iran and Türkiye), and North America (USA). These distributions were incorporated into the analysis in accordance with the Neighbor network tree.

## Acknowledgments

The authors gratefully acknowledge Dr. Yan Zhou, Citrus Research Institute, Southwest University, Chongqing, China, for providing us with the CYVCV coat protein monoclonal antibody 18H5. We also thank Sydney Helm Rodriquez, Biological Science Technician, USDA, ARS, SJVASC, CDPG, Parlier, California, for technical assistance. We are grateful for the support from USDA-APHIS-PPQ-S&T, Raleigh, NC and the CDFA, Sacramento, CA. Mention of trade names or commercial products in this publication is solely for providing specific information and does not imply recommendation or endorsement by the USDA. USDA is an equal opportunity provider and employer.

## Supporting information captions

**Supporting Table 1.** The recombination analysis of *Citrus yellow vein clearing virus* isolates upon genome sequences. Four event were detected in over four different methods implemented in RDP4. Different algorithms abbreviation: R: RDP; G: GENECONV; B: BOOTSCAN; M: MAXCHI; C: CHIMAERA; S: SISCAN; T: 3SEQ.

**Supporting Table 2.** Estimation of selection pressure of *Citrus yellow vein clearing virus*-encoded open reading frames. The positive and negative nucleotide selection sites were also determined by codon-based likelihood algorithms Single-Likelihood Ancestor Counting (SLAC) using Global MG94xREV model with p-value threshold of 0.1. dN/dS >1 indicating positive selection, dN/dS = 1 indicating neutral evolution, and dN/dS <1 indicating negative selection.

**Supporting Fig 1.** A snapshot of the insertion/deletion mutant sites via multiple sequence alignment of 57 *Citrus yellow vein clearing virus* genome sequences. The CYVCV CA isolates were marked with red arrows.

**Supporting Fig 2.** Root-to-tip divergence, as a function of sampling time for Maximum clade credibility tree clusters, of selected *Citrus yellow vein clearing virus* coat protein gene sequences used in this study. Correlation coefficient equals to 0.2457.

**Supporting Fig 3.** *Citrus yellow vein clearing virus* (CYVCV) direct tissue blot immunoassay. No obvious differences were observed between the healthy control and CYVCV-infected Eureka lemon samples. The CYVCV monoclonal antibody 18H5 served as the primary antibody.

**Supporting Fig 4.** The full reconstruction of *Citrus yellow vein clearing virus* (CYVCV) evolutionary history and dynamics of upon Bayesian evolutionary analysis. Colored dots indicate the biogeographic distribution of CYVCV isolates. The circles illustrate the mostly likely distribution status of the most recent common ancestor in key nodes only.

